# Cooperation-based sperm clusters mediate sperm oviduct entry and fertilization

**DOI:** 10.1101/2020.10.18.344275

**Authors:** Yongcun Qu, Qi Chen, Shanshan Guo, Chiyuan Ma, Yonggang Lu, Junchao Shi, Shichao Liu, Tong Zhou, Taichi Noda, Jingjing Qian, Liwen Zhang, Xili Zhu, Xiaohua Lei, Yujing Cao, Wei Li, Wei Li, Nicolas Plachta, Martin M. Matzuk, Masahito Ikawa, Enkui Duan, Ying Zhang, Hongmei Wang

## Abstract

Sperm cooperation has been observed in multiple species, yet its existence and benefit for reproductive success in mammals remains underexplored. Here, combining tissue-clearing with deep three-dimensional imaging, we demonstrate that postcopulatory mouse sperm congregate into unidirectional sperm cooperative clusters at the utero-tubal junction (UTJ), a key physical barrier for passage into the oviduct. Reducing sperm number in male mice by unilateral vasoligation or busulfan-treatment impairs sperm cluster formation and oviduct entry. Interestingly, sperm derived from *Tex101^−/−^* mouse has normal number, motility and morphology, yet they cannot form sperm cluster and fail to pass through the UTJ, which is at least in part due to the altered tail beating pattern of the *Tex101^−/−^* sperm. Moreover, *Tex101^−/−^* sperm’s defect in oviduct entry cannot be rescued by the presence of wild-type (WT) sperm in the same uteri by sequential mating, suggesting sperm cooperative cluster as an essential behavior contributing to male fertility, which could be related to human infertility or subfertility.

## Introduction

Sperm must overcome significant obstacles within the female reproductive track to reach and fertilize the egg. In many mammalian species, the utero-tubal junction (UTJ) is a narrow tube with multiple crypts and viscous fluid(1–4). This structure, connects the uterus to the oviduct and functions as an efficient physical barrier that blocks most sperm in the uterus, allowing only a few to enter the oviduct(5, 6) (Fig 1c). Although this phenomenon has been well-documented for a long time, the mechanisms underlying how sperm passing through the UTJ remain largely unknown.

**Fig 1.**
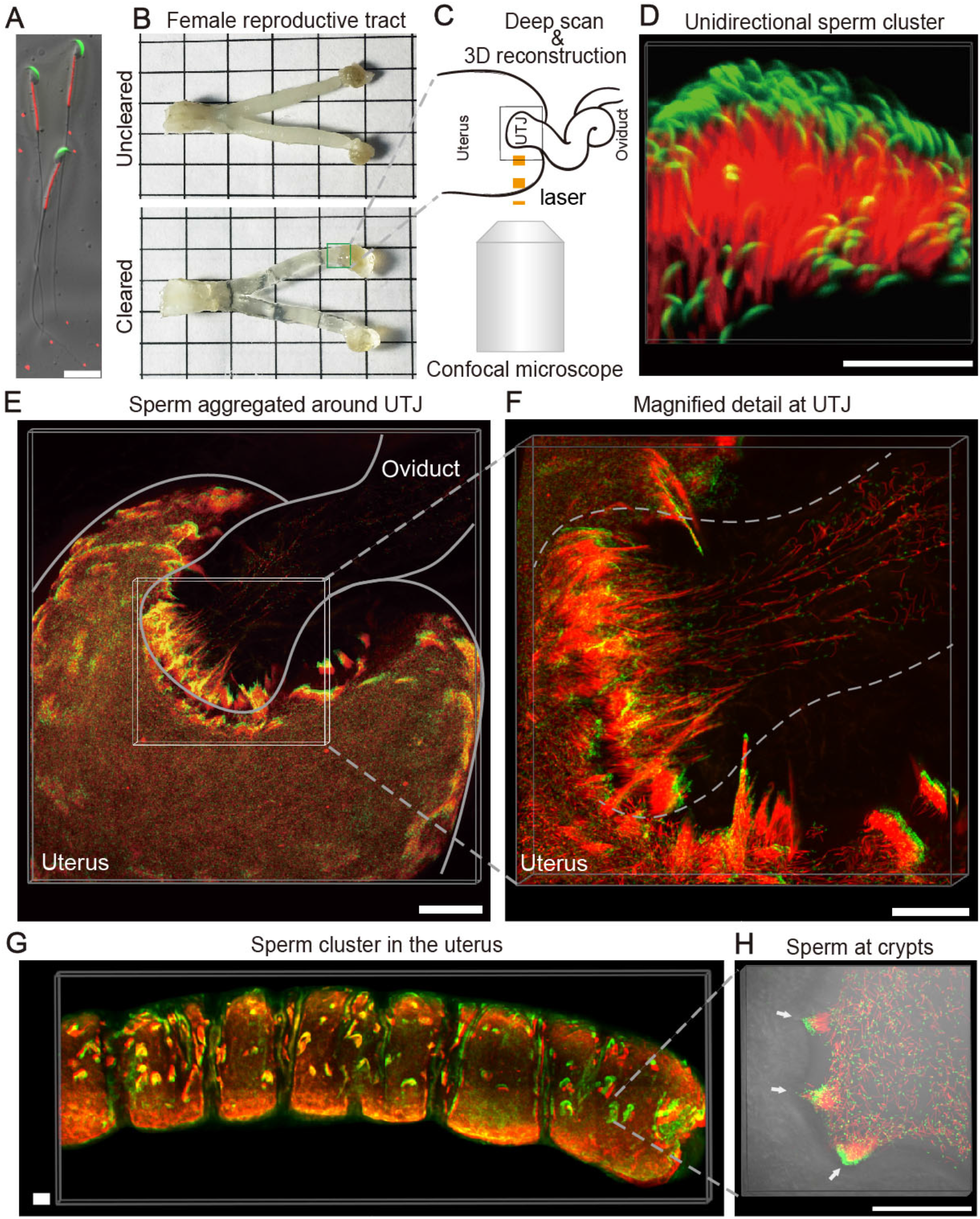
Sperm behavior at the UTJ and in the uterus one hour after coitus. (A) Sperm with GFP-labeled acrosome and RFP-labeled mitochondria (CAG/su9-DsRed2, Acr3-EGFP). Scale bar, 20μm. (B) Female reproductive tract before and after clearing. (C) Schematic of 3D imaging. (D) Sperm cluster with sperm head oriented in the same direction. Scale bar, 30μm (E) 3D imaging of sperm aggregated around the UTJ. Scale bar, 200μm. (F) Magnified detail of sperm cluster at the UTJ. Scale bar, 200μm. (G) 3D imaging of sperm clusters in the uterus. Scale bar, 200μm. (H) Sperm cluster distribution in uterine crypts (white arrow). Scale bar, 200μm.

It has been previously suggested that linear progressive motility can increase the efficiency of sperm to pass through the UTJ(4, 7). However, emerging evidence over past decades indicated that sperm motility alone is not sufficient for sperm oviduct entry(5, 8), as demonstrated by many mouse knockout strains showing normal sperm motility but defected oviduct entry(8, 9), suggesting additional mechanisms. Previously studies in *Drosophila melanogaster* showed that complex sperm behaviors within the female reproductive tract were link to sperm selection and fertilization(10). Recent study in mouse also suggested that additional sperm behavior could be critical for sperm moving through the female reproductive tract(11).

Interestingly, increasing evidences from molluscs to mammals have shown that sperm can group together *in vitro,* a collective behavior which has been termed sperm cooperation, and might be beneficial for mutual advantages to improve fertilization probability(12, 13). For example, several rodent sperm were found to form aggregates in the culture medium to increase progressive motility and to prevent premature acrosome reaction(14, 15). Yet, it remains unclear whether and how mammalian sperm display such behavior *in vivo*, mainly due to the high opacity and inaccessibility of female reproductive tract. Here, by combining tissue clearance, three-dimensional imaging and functional studies, we provide direct evidence showing sperm cooperative cluster at the UTJ; and that the presence of sperm cluster is functionally associated with sperm entry into the oviduct.

## Results

### Sperm aggregate into clusters *in vivo*

Revealing mammalian sperm behavior inside the uterus remains highly challenging due to the opacity of the female reproductive tract. To address this issue, we employed whole-organ clearing technologies, optimized from previous work(16), and this approach rendered the female reproductive tract with high transparency (Fig. 1B). To visualize the intrauterine sperm behaviors at one- and two-hour postcoitum, we used transgenic sperm fluorescently labeled with a red (DsRed2) midpiece, and a green (GFP) acrosome(17) (Fig 1A). Combining with 3-dimensional confocal imaging and reconstruction (Fig 1C), we found that sperm densely assembled into clusters at the UTJ, with sperm heads and tails spatially oriented in the same direction facing the oviduct_(Fig 1D-1F, S1 Fig and S1 Movie). In addition to the sperm clusters found at the UTJ, we found similar clusters inside the uterus, in regions where crypts form (Fig 1G and 1H). These results suggested that sperm clusters may not ‘intentionally’ aggregate at the UTJ, but rather they scout the whole uterus and accumulate at all crypt-like regions, including the UTJ.

### Threshold number for sperm UTJ clustering and oviduct entry

The observation of sperm clusters at the UTJ raised the question whether this population-based sperm behavior is related to sperm oviduct entry. If true, the sperm entry into the oviduct should be hampered when the size of the sperm cluster is reduced. To test this hypothesis, we first generated a unilateral-vasoligated (Uni-Vas) male mice model to approximately reduce the sperm number in half (Fig 2B). Indeed, after two weeks’ recovery, the sperm concentration was significantly decreased to 58.08% of control level (Fig 2D). Using the same method to generate the 3D reconstructed female reproductive tract with sperm, we found that the Uni-Vas males showed a significantly decreased sperm cluster size (volumetric analysis of sperm fluorescent signal) at the UTJ, compared to control mice (Fig 2A, 2B and 2F). Furthermore, the *in vivo* fertilization rate by Uni-Vas males was markedly reduced to 62.62% (control mice: 90.78%) (Fig 2E). These findings demonstrated that the sperm number is a contributing factor to the sperm cluster size at UTJ, and is highly related to the number of sperm that can enter into the oviduct.

**Fig 2.**
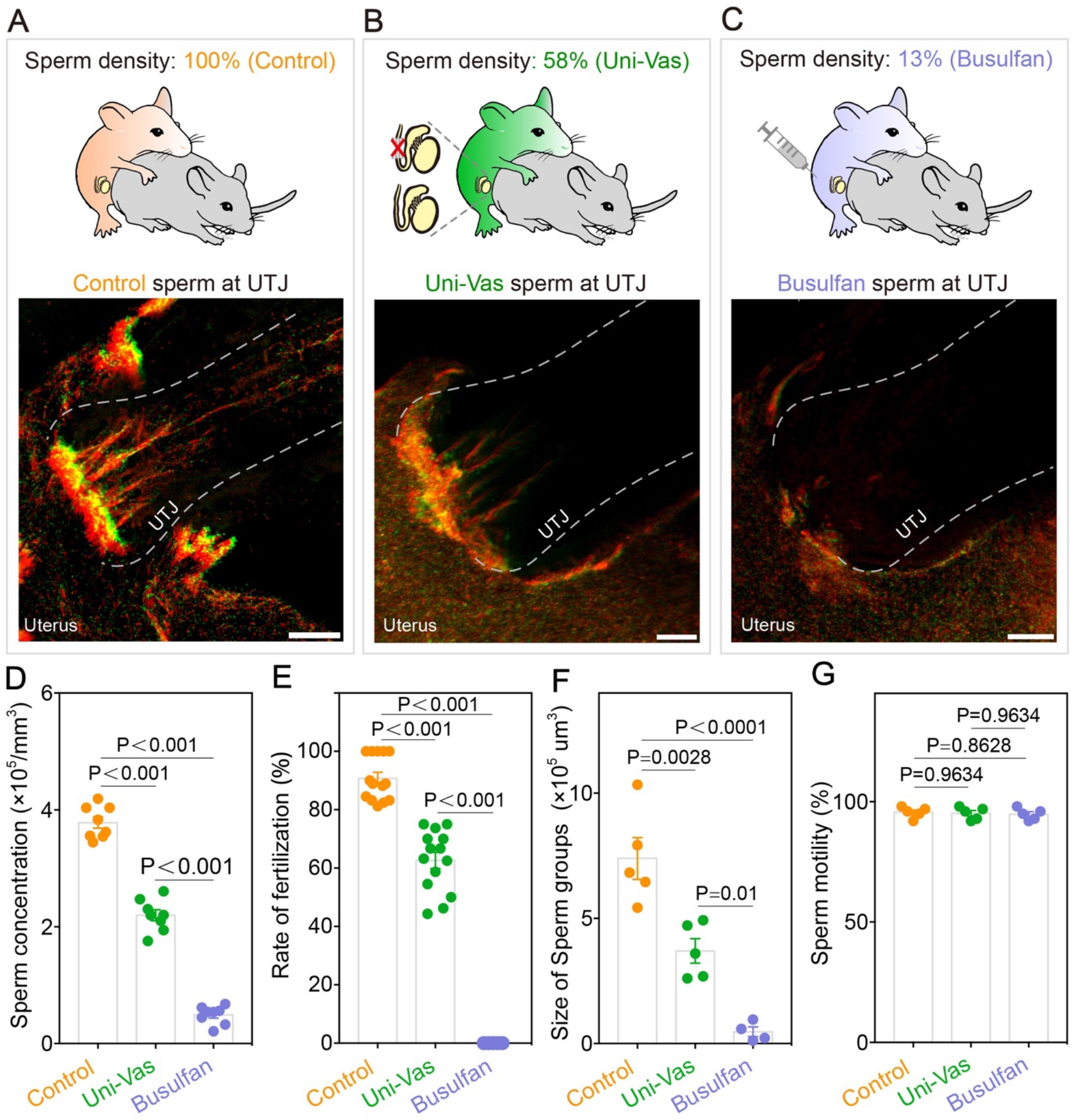
Reduced sperm number decreases the volume of sperm clusters and hampers *in vivo* fertilization. (A-C) Schematic diagrams of mating procedure (top) and 3D visualization of sperm behavior at the UTJ (bottom). Scale bars, 100μm. (D) Analysis of Control, Uni-Vas and Busulfan-treated male sperm concentration in the uteri. n = 8 different visual fields from 4 male mice each. (E) *In vivo* fertilization rate. n = (14 control male mice, 14 Uni-Vas male mice and 5 Busulfan-treated male mice). (F) Quantification of the size of sperm groups (volumetric analysis of the sperm fluorescent signal) in control, Uni-Vas and Busulfan-treated male at the UTJ (two hours after coitus). n = (5 control male mice, 5 Uni-Vas male mice and 4 Busulfan-treated male mice). (G) sperm motility of Control, Uni-Vas and Busulfan-treated male mice. n=5 male mice each. The results are shown as mean ± SEM.

To further reduce the sperm number, we used another male mouse model by injecting a single dose of busulfan (Fig 2C), a cytotoxic regent that can kill spermatogonia and induce oligospermia or azoospermia. Two months after a single low-dose (17mg/kg) busulfan treatment, the sperm concentration reduced to 5.65%-17.9%(from 5 mice, higest:17.90% and lowest: 5.65%) of control level (Fig 2D), causing a complete infertility in these mice (Fig 2E), although the motility of the remaining sperm showed no difference compared with control mice (Fig 2G). Importantly, 3D imaging results showed that despite the fact that the sperm number decreased substantially, some sperm could still reach the UTJ, but in this situation, little or no sperm cluster could form at the UTJ (Fig 2C and 2F) and no sperm could enter the oviduct (Fig 2E). These results further supported the conclusion that a sufficient number of sperm is essential for the efficient formation of sperm cluster at UTJ, which is highly related to sperm’s ability passing through the UTJ. When the sperm number decrease below a certain threshold, the sperm cannot pass through the UTJ. In addition, according to our mating data, we found that when sperm concentration was close to 25% of pretreatment level, sperm could be found in the oviduct and the mice showed subfertility (~22% fertility rate compared to the control mice). Thus, we deduced that the minimal of sperm counts that can support the sperm entrance into the oviduct could be between 17.9% to 25% of control level. This finding was consistent with previous evidence suggesting that ~20% of normal sperm counts is a threshold for male mice fecundity(18).

### Sperm clustering is controlled by genetic factors

In addition to the importance of the sperm counts for sperm clustering and fertility, other factors should be also involved. Indeed, recent studies have reported that over ten different knockout mouse models that showing normal sperm counts, motility, and morphology, but the sperm from these mutant mice cannot migrate through UTJ and these male mice are infertile(8, 9). Could these defects relate to sperm clustering? To test this hypothesis, we used one of these knockout strains, *Tex101*^−/−^(19, 20) to test the sperm ability to aggregate *in vitro* (Fig 3) and to form cluster at UTJ *in vivo* (Fig 4).

**Fig 3.**
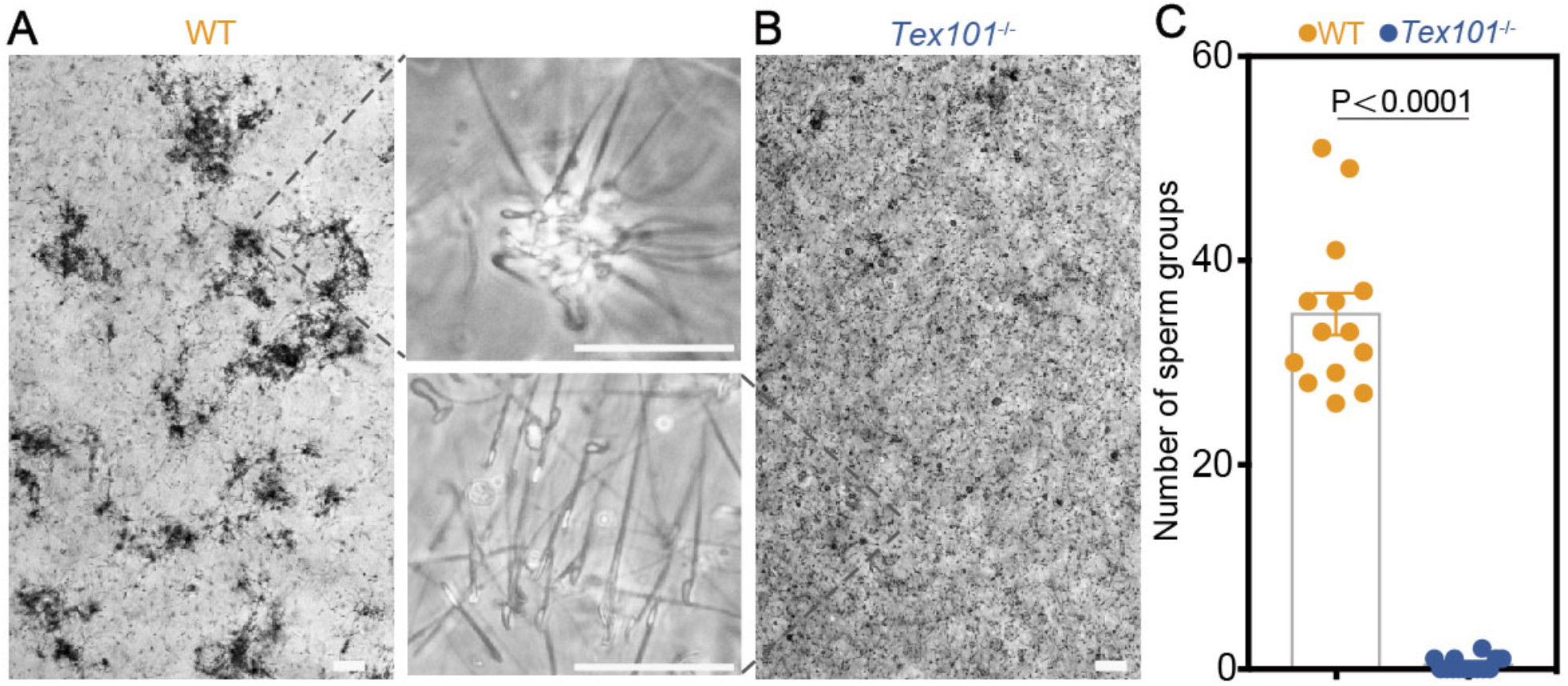
Comparison of WT and *Tex101*^−/−^ aggregation ability *in vitro.* (A) WT sperm aggregate during *in vitro* culture. (B) *Tex101*^−/−^ sperm show impaired aggregation. (C) Quantification of aggregated sperm groups (> 10 sperm per group) for WT and *Tex101*^−/−^ mice. The results are shown as the mean ± SEM, n = 14 different visual fields from 4 male mice each. Scale bars, 50μm.

**Fig 4.**
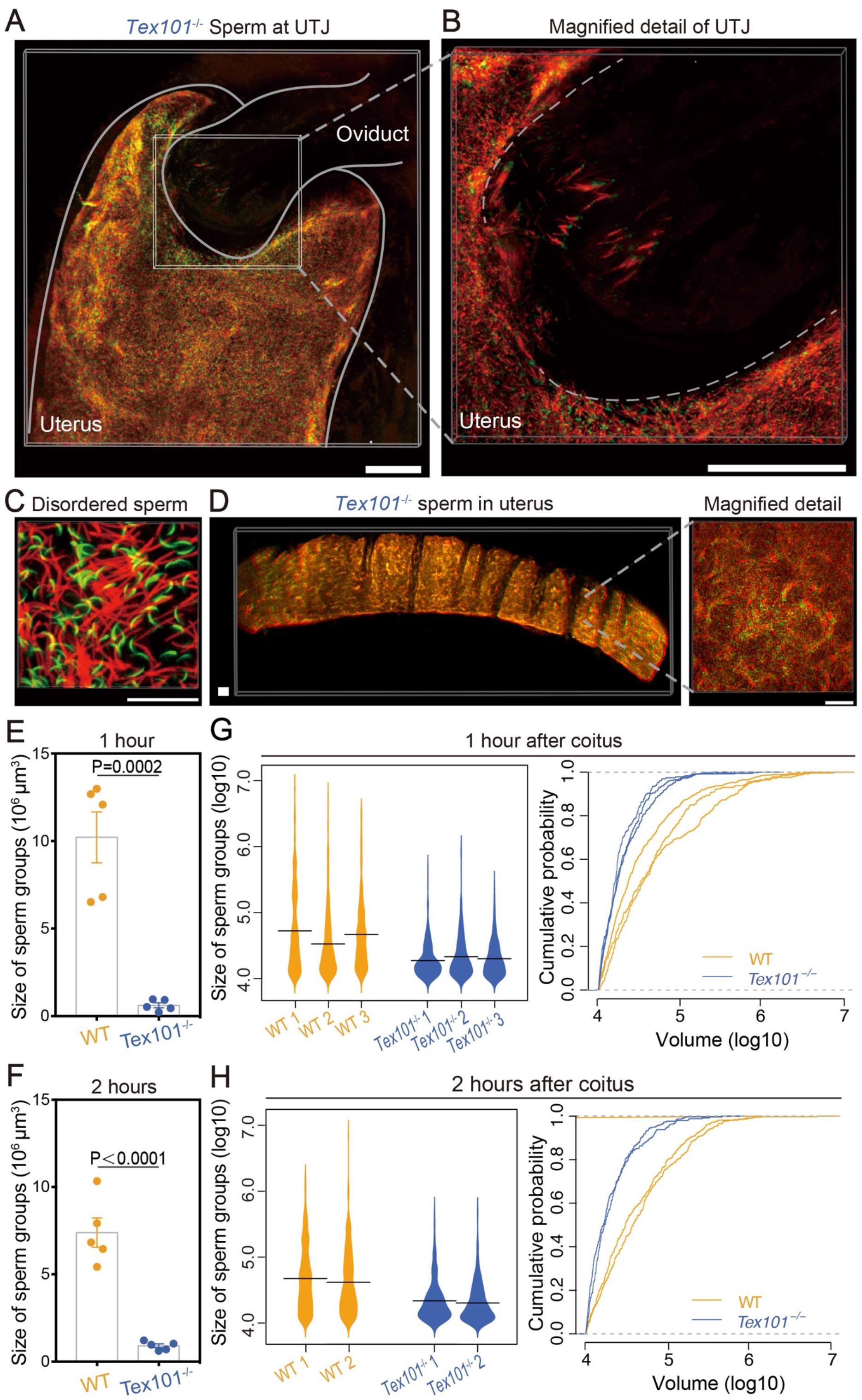
*Tex101*^−/−^ Sperm behavior at the UTJ and in uterus. (A) *Tex101*^−/−^ sperm fail to form clusters at the UTJ. Scale bar, 200μm. (B) Magnified detail of *Tex101*^−/−^ sperm at UTJ. Scale bar, 200μm. (C) *Tex101*^−/−^ sperm displaying disordered distribution. Scale bar, 30μm. (D) *Tex101*^−/−^ sperm remain scattered in the uterus. Scale bars, 200μm. (E-F) Quantification of the size of sperm groups (volumetric analysis of the sperm fluorescent signal) in WT and *Tex101*^−/−^ at the UTJ at one (E) and two hours (F) after coitus on a red fluorescence signal-generating surface. (G-H) Quantitative comparison of the size of sperm groups (volumetric analysis of the sperm fluorescent signal) for the WT sperm aggregated into sperm clusters and the scattered *Tex101*^−/−^ at one (three WT and three *Tex101*^−/−^ replicates) (G) and two hours (two WT and two *Tex101*^−/−^ replicates) (H) after coitus, respectively. Left panel is a violin plot and the right panel is the cumulative distribution. Results are shown as the mean ± SEM, n = 5 male mice each in (E) and (F), n=3 male mice in (G), n=2 male mice in (H).

For the *in vitro* aggregation experiment, cauda epididymal sperm of WT and *Tex101*^−/−^ were separately released into the M16 medium and then the numbers of the sperm aggregation (> 10 sperm per aggregation) between these two groups were compared. As shown in Fig 3, the aggregation ability of *Tex101*^−/−^ sperm was severely damaged. Similar defects of *in vitro* aggregation has also been observed in other gene knockout mice that showing normal sperm counts but with infertile phenotype(21).

To further exam whether the *Tex101*^−/−^ sperm can form sperm cluster *in vivo,* we crossed the *Tex101*^−/−^ mice into the same transgenic mice (red (DsRed2) midpiece, and green (GFP) acrosome), mated these mutant mice with normal female, and then used the same tissue-clearing and 3D imaging analyses as we did for WT sperm. As shown in Fig.4, the 3D imaging showed that the sperm of *Tex101*^−/−^ mice were unable to form cluster *in vivo* (Fig 4A, 4B, 4E, 4F and S2 Fig); instead, they distributed irregularly near the UTJ (Fig 4C), and most of them were blocked outside or at the entry of the UTJ (Fig 4B and S2 Movie).

Moreover, according to the volumetric analysis of the sperm groups (depending on the fluorescent signal) in the uterus, we also found that *Tex101*^−/−^ sperm could not form large clusters within the uterus, but only have tiny groups with disordered orientation compared with the WT mice (Fig 4D and 4G and 4H). These results supported the idea that in addition to sperm counts, morphology and motility, the sperm clustering behavior could be another essential contributing factor for efficient sperm migration and fertility.

### Sequential mating experiments demonstrate that sperm clustering is an intrinsic property independent of sperm number

Subsequently, we tested whether those sperm that cannot form sperm cluster, such as those derived from the *Tex101*^−/−^ mouse, would be expected to enter the oviduct with the help of a normal sperm cluster. To test this, we developed a mating procedure allowing a wild-type female to sequentially mated with two transgenic males bearing sperm with different markers. This sequential mating experiment is possible as mice are promiscuous(22, 23). We used WT (GFP-labeled sperm tail)(24) followed by *Tex101*^−/−^ (RFP-labeled sperm tail)(17), and *Tex101*^−/−^ (RFP-labeled sperm tail) followed by WT (GFP-labeled sperm tail) (Fig 5A and 5C). Video monitoring confirmed that successful mating with the second male occurred as early as 45 min after the first male. Imaging of WT and *Tex101*^−/−^ sperm in the uterine horns after sequential mating revealed that only WT sperm formed clusters and entered the oviduct. By contrast, *Tex101*^−/−^ sperm could not migrate into the oviduct, albeit they entered the uterus before the WT sperm (Fig 5B and 5D). The imaging data were further supported by functional results, showing that after sequential mating, all offspring derived from WT and none from *Tex101*^−/−^ sperm, independently of mating order (Table 1). Therefore, although the sperm number in the uterus increased substantially after sequential mating, the scattered *Tex101*^−/−^ sperm cannot either form cluster or take advantage of the sperm cluster of WT sperm to enter the oviduct, indicating the sperm clustering itself is an intrinsic property responsible for sperm entry into the oviduct.

**Fig 5.**
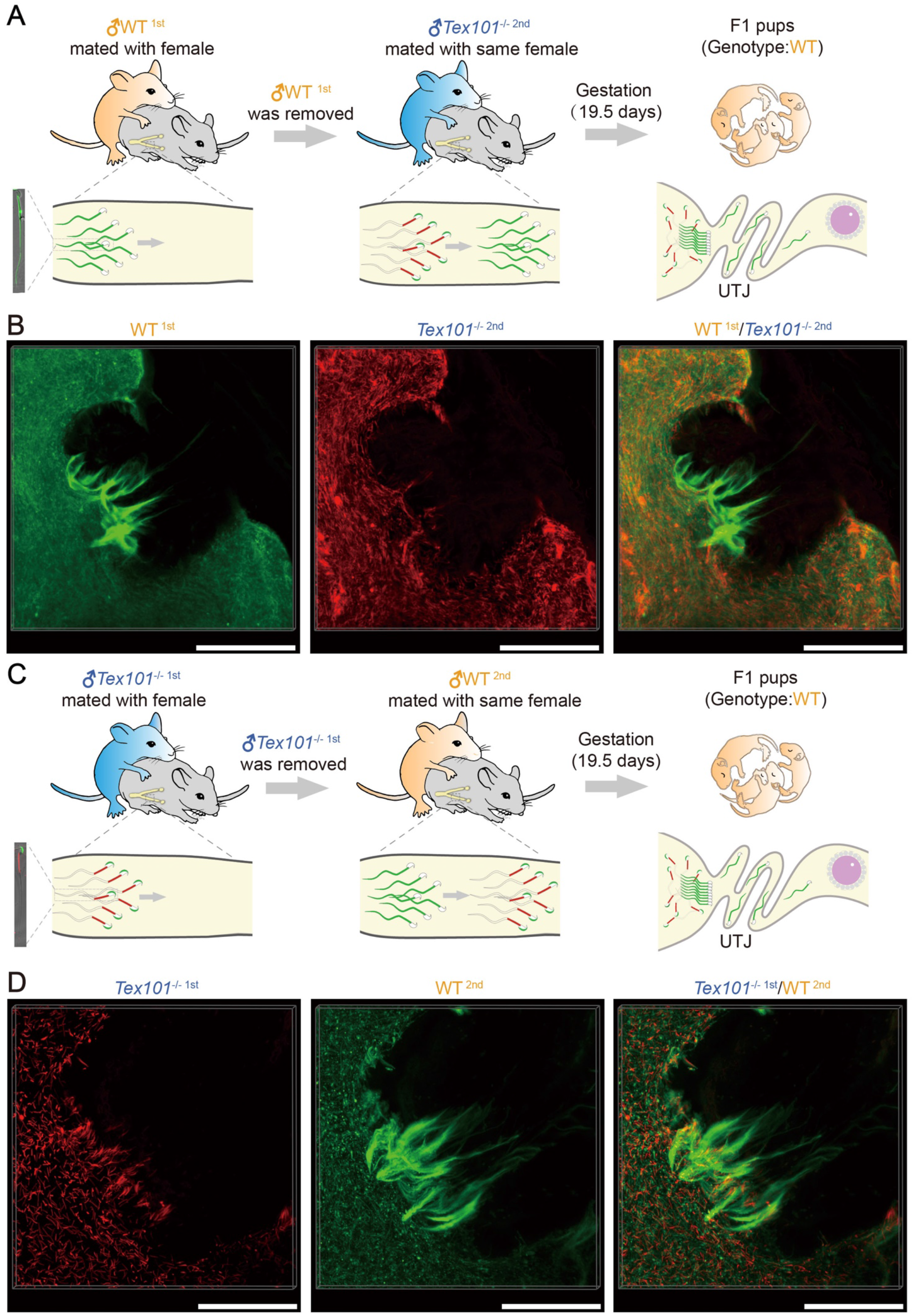
Sequential mating experiments showing that sperm clustering is necessary for passing through the UTJ. (A, C) Schematic diagrams of sequential mating experiment. In (A) the female mated first with the WT male and then with the *Tex101*^−/−^ male (WT^1st^, *Tex101*^−/−2nd^). In (C) the female mated first with the *Tex101*^−/−^ male and then with the WT male *Tex101*^−/−1st^, WT^2nd^). (B, D) 3D visualization of fluorescently labeled WT and *Tex101*^−/−^ sperm behavior at the UTJ after sequential mating. In (B) the female mated first with the WT male and then with the *Tex101*^−/−^ male (WT^1st^, *Tex101*^−/−2nd^). In (D) the female mated first with the *Tex101*^−/−^ male and then with the WT male (*Tex101*^−/−1st^, WT^2nd^). Scale bars, 200μm.

**Table 1.** Litter size and genotype after sequential mating.

|  |  | Number of pups |  |  |
| --- | --- | --- | --- | --- |
| Mating order | Number of cages | Litter size | WT | *Tex101*<sup>-/-</sup> |
| WT<sup>1st</sup><br>*Tex101*<sup>-/-</sup> 2nd | 1 | 19 | 19/19 | 0/19 |
|  | 2 | 9 | 9/9 | 0/9 |
|  | 3 | 11 | 11/11 | 0/11 |
|  | 4 | 5 | 5/5 | 0/5 |
|  | 5 | 13 | 13/13 | 0/13 |
|  | 6 | 12 | 12/12 | 0/12 |
|  | 7 | 6 | 6/6 | 0/6 |
|  | 8 | 9 | 9/9 | 0/9 |
| *Tex101*<sup>-/-</sup> 1st<br>WT<sup>2nd</sup> | 1 | 12 | 12/12 | 0/12 |
|  | 2 | 14 | 14/14 | 0/14 |
|  | 3 | 7 | 7/7 | 0/7 |
|  | 4 | 8 | 8/8 | 0/8 |
|  | 5 | 9 | 9/9 | 0/9 |
|  | 6 | 5 | 5/5 | 0/5 |
|  | 7 | 10 | 10/10 | 0/10 |
|  | 8 | 11 | 11/11 | 0/11 |

### Impaired clustering in *Tex101*^−/−^ sperm is associated with altered sperm tail beating pattern

Why the *Tex101*^−/−^ sperm cannot for clusters? One possibility is that this could be due to that the sperm lack certain “sticky” molecules as we previously discussed(19, 20). In the currently study, we further explored an alternative explanation that the lack of grouping behavior might be related to abnormal sperm swimming pattern and altered hydrodynamics, as there is a recent study showing that sperm collective behavior formation is influenced by the sperm waveform dynamics(25). To test this hypothesis, we released sperm (WT vs *Tex101*^−/−^) from uteri 1.5 hours after coitus (mating with normal C57 female) and put the sperm into the viscoelastic medium, followed by detailed video recording/analysis of sperm swimming in a frame-by-frame manner (Fig.6). This detailed analysis has led to an exciting discovery that the WT and *Tex101^−/−^* sperm indeed showed different swimming patterns. As shown in Fig 6A and S3 movie, the sperm flagellar bending pattern of anti-hook and pro-hook were almost symmetric distributed in WT sperm. However, the *Tex101*^−/−^ sperm have two asymmetric flagellar bending patterns, showing a prolonged pro-hook bending and a brief anti-hook bending besides the pro-hook bending (Fig 6B and S4 Movie), the extent of both pro-hook bending and a brief anti-hook (reflected by the minimal value of angle θ in Fig 6A and 6B) in *Tex101*^−/−^ sperm were more dramatic than the WT sperm (Fig 6C-F). This different swimming pattern might hold the key to explain why *Tex101*^−/−^ sperm cannot form cluster, because this more dramatic asymmetric anti-hook/pro-hook swimming pattern might make the sperm head difficult to “attach” to the objectives. This may partially explain why the *Tex101*^−/−^ sperm cannot group at the uterine crypts or the UTJ.

**Fig 6.**
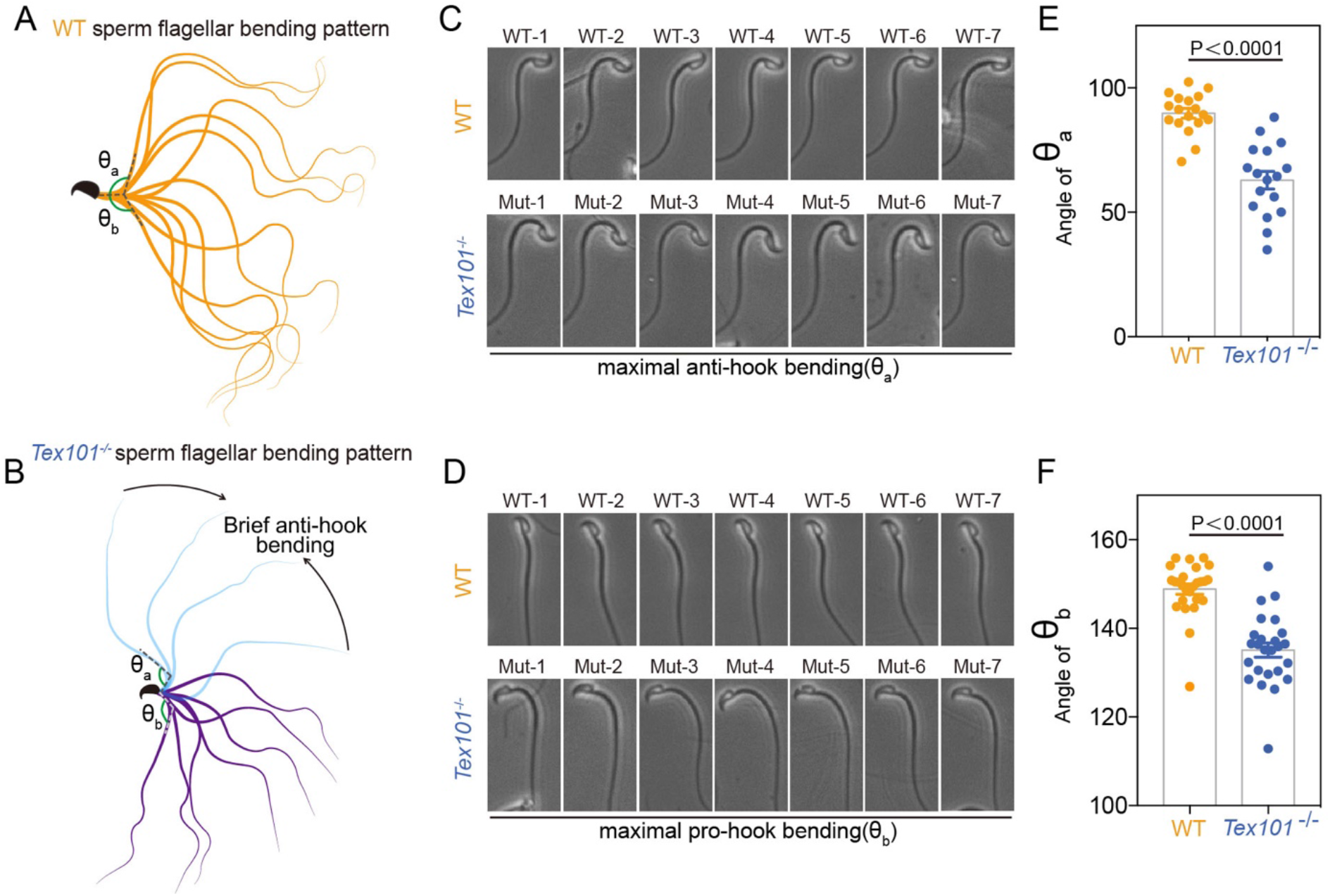
Comparison of WT and *Tex101*^−/−^ sperm flagellar bending pattern. (A) WT sperm flagellar bending pattern. (B) *Tex101*^−/−^ sperm flagellar bending pattern. (C) Images of WT (top) and *Tex101*^−/−^ (bottom) sperm maximal anti-hook bending. (D) Images of WT (top) and *Tex101*^−/−^ (bottom) sperm maximal pro-hook bending. (E) Angle of WT and *Tex101*^−/−^ sperm maximal anti-hook bending around the neck. (F) Angle of WT and *Tex101*^−/−^ sperm maximal pro-hook bending around the neck. The results are shown as the mean ± SEM, n = 4 male mice each.

Moreover, during the detailed recording on the sperm that are freshly released from the uteri (1.5 hours after coitus), we have obtained further insight on how the sperm might change their moving pattern after they form cluster. In S3A Fig and S5 movie, we found that when a cluster of sperm swimming around cell debris, all the sperm tails wave in a synchronized manner, generating a strong tail pendulum force pushing the cell/debris. Since the sperm clusters at the UTJ were positioned in a unified direction towards the UTJ (Fig 1D-1F), this very force generated by the synchronized sperm tail beating might push the UTJ open and allow the sperm at the center of the entrance to penetrate into the oviduct. Whereas the *Tex101*^−/−^ sperm could not form the cluster around cell/debris and form this unified force (S3B Fig and S6 Movie). This data obtained from ex vivo sperm video could provide explanations on two facts: **1)** a reduced sperm number, either in the Uni-vas or busalfan treatment experiments (Fig. 2), decreases the overall force generated by the synchronized sperm tail beating, and when the generated force is below a certain threshold it cannot efficiently push the UTJ open. **2)** The *Tex101*^−/−^ sperm never achieve the threshold of force to push open the UTJ because it cannot form sperm cluster at UTJ (Fig 4).

## Discussion

In the present work, we demonstrate that mouse sperm establish unidirectional sperm clusters at the UTJ, and that this intrauterine sperm cooperation-based behavior contribute to sperm oviduct entry and fertility. Although *in vitro* experiments had revealed that mouse sperm aggregation might be related to male fertility(21), it remains an unsolved issue whether this sperm cooperation behavior exist within the female tract after coitus. Traditional histological methods applied on thin tissue sections provide limited spatial information about sperm behavior *in vivo.* We addressed this technical challenge by combining optical sectioning microscopy and optimized tissue clearing techniques(26). As the post-copulation uterus contains liquid semen, we used hydrogel-based tissue clearing(16) to render the post-copulation female reproductive tract with copulation plug transparent. This approach improved semen fluid density and turned uterine contents into an elastic gel, which enabled the fixation of large sperm population and the preservation their position as they were *in vivo.* Using this method, we successfully discovered the sperm cooperation behavior in the female reproductive tract after mating.

Why do sperm aggregate to form groups when they swim through the female reproductive tract? From evolutional perspective, sperm evolution is mainly drives by two forces: postcopulatory sexual selection from female cryptic choice and sperm competition environment between males(27, 28). From the female perspective, making all her eggs fertilized by health sperm is the priority, thus female reproductive tract designs complex barriers to select sperm(5). but from the male part, all the sperm are potentially valuable and could be in some way to facilitate fertilization to overcome the female “obstacles” and to engage with competitor’s sperm in polyandrous environment(12). In this situation, male may evolve sperm cooperation behavior to efficiently transport through the female tract and response to dramatic sperm competition environment(29). Experimental studies have found that the coitus induces a viscous oviduct fluid flow towards the UTJ, which is thought to reduce sperm beat frequency and hinder their migration(30). Our current study has provided further compelling evidence that sperm clustering *in vivo* is functionally important as the sperm cluster can provide enhanced force to push the UTJ open, thus enable the sperm entrance into the oviduct. When the sperm cannot form cluster, such as in the case of *Tex101*^−/−^ mice, the sperm are blocked outside the narrow UTJ. Interestingly, we also found that sperm predominately aggregate into clusters in uterine crypts, which are anatomical structure similar to the UTJ. Although lacking of exact evidence, we speculated that sperm cells might find their way to the UTJ via a trial-and-error process, and the mechanism of sperm cluster formation may also involve in sperm-female and sperm-sperm communication(31).

The molecular and physiological mechanisms by which sperm congregate into the sperm cluster within the intrauterine environment remain unknown. Plausibly, they may rely on the cohesive nature of sperm, which is lost in some gene knockout mice(21), for example, *Tex101*^−/−^ mutant. Additionally, we found that the sperm movement pattern may also contribute to the sperm cluster formation (Fig 6), which resonates with previous report that collective behavior could be driven by hydrodynamic interactions regulated by sperm waveform dynamics(25). Following ejaculation, sperm are in close proximity with each other at a high concentration, which could facilitate sperm-sperm interactions and generate synchronized tail beating to enhance the sperm force to push objects they attached to (e.g. the UTJ). Furthermore, the highly viscous fluid within the female reproductive tract with low Reynolds number(32), may hydrodynamically promote rapid sperm clustering and synchronized swimming, where decreased sperm density and abnormal beating wavelength may disrupt sperm clustering(25, 33). Moreover, the confined geometry of the convoluted epithelial lining may also contribute to sperm clustering, as most clusters aggregate within uterine crypts and UTJ. Previous studies have suggested sperm has strong corner-swimming behavior, which was originate from hydrodynamic interaction of sperm with surface of the corner(34). Notably, similar population-based sperm behavior may also exist in human, and it has been reported that high density of sperm was located at crypts of human cervix(35), where the sperm may similarly form cluster and generated synchronized force to facilitate their moving into uterus.

Another important finding of our work is that decreased sperm number hampered sperm cluster formation at the UTJ and sperm oviduct entry. According to our data, despite having normal morphology and motility, when sperm counts drop to 17.9% (or less) of control level, sperm cluster is hard to form and none of the sperm can enter the oviduct.

Finally, our study is also highly related to human fertility. While sperm counts, motility, and morphology are clinically recognized as indicators of male fecundity, it has been suggested that these factors often cannot fully account for or predict clinical diagnosis(36). Defects in sperm clustering may account for certain infertility or subfertility problems in human with normal sperm count and morphology. Our data may also explain subfertility in oligospermia (low sperm counts), which could be due to insufficient sperm clustering in the female reproductive tract thus fail to migrate through major barriers such as the cervix. In conclusion, our study revealed that the sperm cooperation behavior takes part in the process of fertilization and link to the male fertility, which would provide a new insight for the research in evolutionary and reproductive biology.

## Material and Methods

### Ethics statements

All animal experiments were conducted under the protocol and animal ethics guidelines of the Animal Care and Use Committee of Institute of Zoology, Chinese Academy of Sciences.

### Mice

Seven-week-old virgin female and adult male mice (C57BL/6) were purchased from Charles River Laboratories China Inc. *Tex101*^−/−^ female mice were mated with *[B6D2F1-Tg-(CAG/su9-DsRed2, Acr3-EGFP) RBGSOO2Osb]* male mice to generate *Tex101*^+/-^ transgenic mice and then backcrossed to generate *Tex101*^−/−^-[*B6D2F1-Tg-(CAG/su9-DsRed2, Acr3-EGFP) RBGSOO2Osb*]. Adult WT-[*B6D2F1-Tg-(CAG/su9-DsRed2, Acr3-EGFP) RBGSOO2Osb*] male mice were unilaterally vasoligated (Uni-Vas) after anesthetized by intra-peritoneal administration of ‘Avertin’ (300mgkg^-1^ body weight) and used for *in vivo* fertilization detection and imaging after resting for two weeks. Adult WT-[*B6D2F1-Tg-(CAG/su9-DsRed2, Acr3-EGFP) RBGSOO2Osb*] male mice were injected a single low-dose(17mg/kg) busulfan and rest for six weeks, after that, used for *in vivo* fertilization detection and imaging once a week. WT-Ddx4-cre(T/W);mT/mG(mut/wt) male mice (GFP-labeled sperm tail) were used for sequential mating. Mice were maintained with a 12:12-hour light-dark cycle, and the health status is specific pathogen-free according to the Animal Care and Use Committee of Institute of Zoology, Chinese Academy of Sciences.

### Conventional mating and female reproductive tract collection

*WT-[B6D2F1-Tg-(CAG/su9-DsRed2, Acr3-EGFP)RBGS002Osb]* male mice, *Uni-Vas-[B6D2F1-Tg-(CAG/su9-DsRed2, Acr3-EGFP)RBGS002Osb]* male mice, Busulfan-treated-[*B6D2F1-Tg-(CAG/su9-DsRed2, Acr3-EGFP)RBGS002Osb*] male mice, and *Tex101*^−/−^-[*B6D2F1-Tg-(CAG/su9-DsRed2, Acr3-EGFP)RBGS002Osb*] male mice were mated with superovulated female mice after human chorionic gonadotropin (hCG) injection. Ejaculation was verified by male shiver followed by a motion that the female was immobilized by the male(22). The time of the appearance of the copulation plug was recorded as 0 hours postcoitum. The female mice were sacrificed by cervical dislocation and the whole female reproductive tract with the inner contents was collected with the copulation plug at 1 hour or 2 hours after coitus and placed into 4% paraformaldehyde (PFA, Sigma) at 4°C overnight.

### Sequential mating and sample collection

In the first group, superovulated female mice mated with WT-Ddx4-cre(T/W);mT/mG(mut/wt) male mice whose sperm are labeled with GFP, and the mice were separated immediately after successful ejaculation. The postcoitum female mice continued to mate with *Tex101*^−/−^*-[B6D2F1-Tg-(CAG/su9-DsRed2, Acr3-EGFP)RBGS002Osb])* male mice (WT^1st^/*Tex101^−/−2nd^*). In the second group, the mating order of the male was switched *Tex101^−/−1st^/WT^2nd^).* Female mice with two successful copulations were sacrificed by cervical dislocation and used for imaging and litter size detection, as we collected the reproductive tracts with inner contents from these females 1 hour after the second copulation for imaging and the other female mice were caged alone to record their litter sizes separately. Offspring were genotyped by PCR at the end of pregnancy.

### *In vivo* fertilization

WT-[*B6D2F1-Tg-(CAG/su9-DsRed2, Acr3-EGFP)RBGS002Osb]* male mice, Uni-Vas-[*B6D2F1-Tg-(CAG/su9-DsRed2, Acr3-EGFP)RBGS002Osb]* male mice, Busulfan-treated-[*B6D2F1-Tg-(CAG/su9-DsRed2, Acr3-EGFP)RBGS002Osb]* male mice, and *Tex101*^−/−^-[*B6D2F1-Tg-(CAG/su9-DsRed2, Acr3-EGFP)RBGS002Osb*] male mice respectively were mated with virgin female mice after hCG injection. Twenty-five hours later, the female mice were sacrificed by cervical dislocation and the embryos were collected from the oviducts. The fertilization rate was detected by the ratio of fertilized zygotes (with two pronuclei) compared with total embryos.

### Whole-organ clearing procedure

An aqueous-based technique named PACT (*pa*ssive *c*larity *t*echnique) was optimized applied for whole-organ clearing as previously reported(16). Briefly, PFA-fixed whole female reproductive tracts were washed with 1 ×PBS 3-4 times and then immersed in 8% of (29% acrylamide: 1 % bisacrylamide solution) mixed with 0.25% VA-044 initiator (Wako Chemicals) at 4°C on nutator for 24 hours. The immersed female reproductive tracts were degassed and hybridized at 37°C for 3 hours. Samples were cleaned up with excessive hydrogel on the surface and then placed into 8% SDS at 37°C with shaking for 3 days. After 3 days of SDS incubation, the female reproductive tracts were washed with 1 ×PBS 5-6 times for 12 hours at 37°C with shaking. Then, the samples were transferred to a concentration of 88% Histodenz w/v (Sigma) for 3 days.

### Image data obtaining and analysis

Clarified female reproductive tracts were placed on glass bottom culture dishes (NEST) and imaged with a Zeiss LSM 780 confocal microscope using 10X and 25X lenses. Imaging data were analyzed with Imaris software (8.4.1, Bitplane). Tile scans were stitched using Zeiss software. The RFP signal was used to generate a surface under 3D surpass mode for quantitative analysis. The surface gain size and diameter of the largest sphere were fit into the object, and the threshold value setting was consistent with image signal. Total relative sperm volume was obtained based on RFP signal (on sperm tail) under 3D surface module, and then sperm concentration was determined by *per unit volume of total sperm/per unit volume of uterine lumen.*

### Sperm motility analyses

Male mice were sacrificed by cervical dislocation and sperm obtained from the cauda epididymis was performed by directly releasing into the TYH medium. After 15 min incubation, sperm at 37C° with a Slide Warmer (# 720230, Hamilton Thorne Research) was detected motility by CASA system (Version.12 CEROS, Hamilton Thorne Research).

### *In vitro* sperm aggregation experiment

Male mice were sacrificed by cervical dislocation and sperm were obtained from the cauda epididymis of 8- to 10-week-old WT and *Tex101*^−/−^ male mice, and the sperm aggregation experiment was processed as previously described(21). Briefly, sperm were released into M16 medium (Sigma) containing 0.5% bovine serum albumin (Sigma), and sperm concentration was adjusted to 2-4 × 10^6^μm^3^. After incubation at 37°C/5% CO2 for 30 minutes, aggregated sperm groups (at least > 10 sperm cells) were observed using a Nikon ECLIPSE TS100 microscope.

### Sperm flagellar bending pattern analysis

Female mice successfully mated with WT or *Tex101*^−/−^ male mice were sacrificed by cervical dislocation 1.5h after coitus. Sperm in the uterine horn were directly released into the preheated viscoelastic medium (0.7% LC-PAM in 0.1M PBS), and then putting into the sperm analysis chamber (Beverly, depth: 20 μm) at 37*C* with a Slide Warmer. Sperm flagellar bending patterns were recorded by NIS-BR software under Nikon ECLIPSE Ti microscope. Sperm maximal anti-hook and pro-hook bending angles around the neck were determined by Image J.

### Sperm swimming pattern around the cell debris obtaining

Female mice successfully mated with WT or *Tex101*^−/−^ male mice were sacrificed by cervical dislocation 1.5h after coitus. Semen obtained from the uterine horn were added into sperm analysis chamber (Beverly, depth:20μm) at 37C with a Slide Warmer. Sperm movement patterns around the cell debris were recorded by NIS-BR software under Nikon ECLIPSE Ti microscope.

### Data statistics

Violin plot and cumulative distribution methods were used to analyze sperm group distributions based on the relative size of sperm groups (volumetric analysis of fluorescent signal) in utero (Twotailed Kolmogorov-Smirnov test). Other quantitative data were subjected to analysis by a two-tailed unpaired Student’s t-test (2 groups), or an ordinary one-way ANOVA (>2 groups) with multiple comparisons test. The results were analyzed with GraphPad Prism 7 and presented as mean ± standard error (SEM); statistical significance was set as P < 0.05.

## Acknowledgments

This work was supported by the National Key Research and Development Program of China (2019 YFA0802600 to YZ, 2018YFC1004500 to YZ, 2016YFA0500903 to ED, 2017YFC1001401 to ED, 2015CB943003 to YZ), the Strategic Priority Research Program of the CAS (XDA16020700 to HW) National Basic Research Program of China (81490742 to ED), National Natural Science Foundation of China (81490741 to HW, 31671201 to YZ, 31671568 to ED), Youth Innovation Promotion Association, CAS (Grant No. 2016081 to YZ), NIH (R01HD092431 to QC). We thank Shiwen Li for the help of image processing.

## Supporting information

**S1 Fig.**
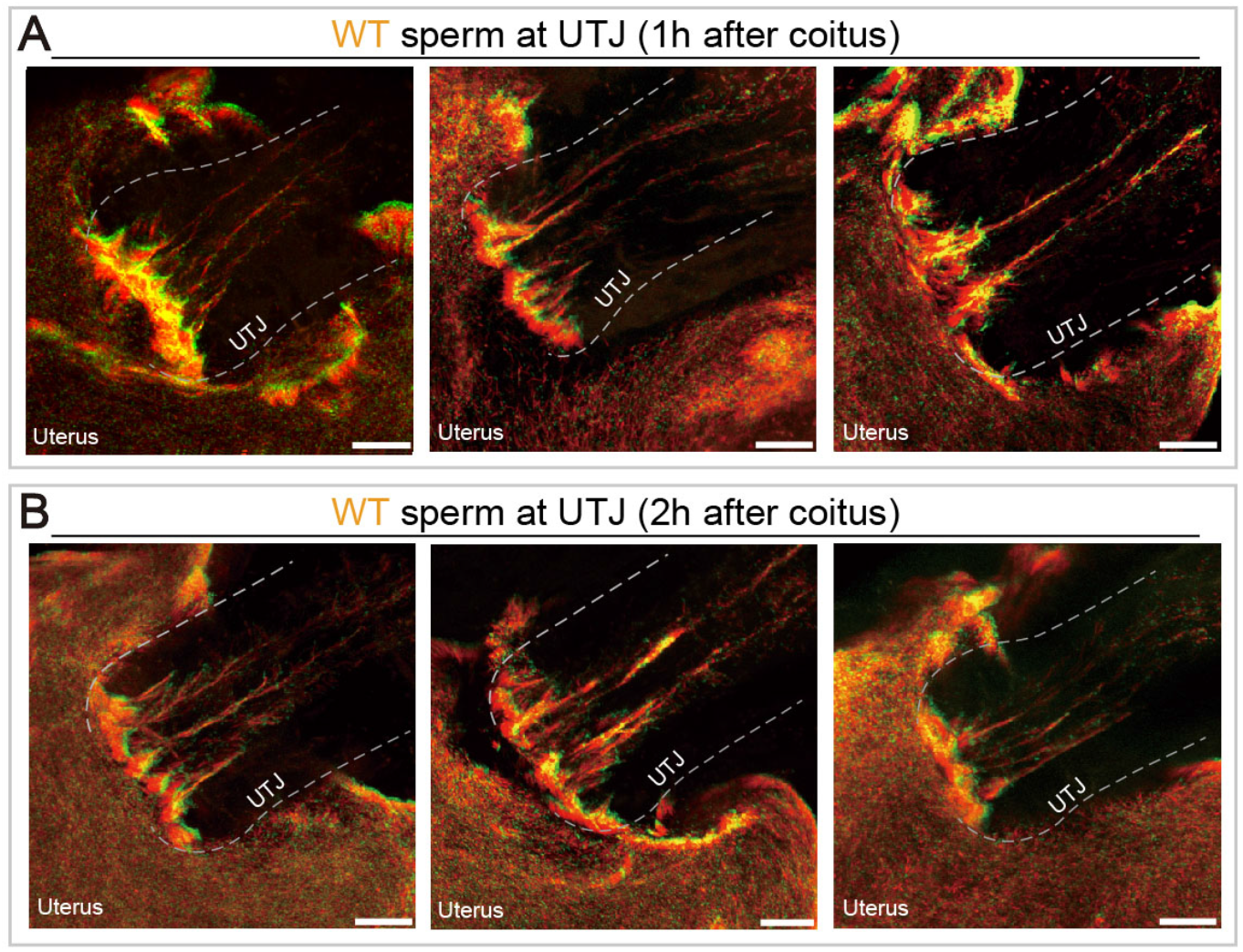
WT sperm behavior at UTJ. (A) WT sperm behavior at UTJ 1h after coitus. (B) WT sperm behavior at UTJ 2h after coitus. scale bars, 200μm.

**S2 Fig.**
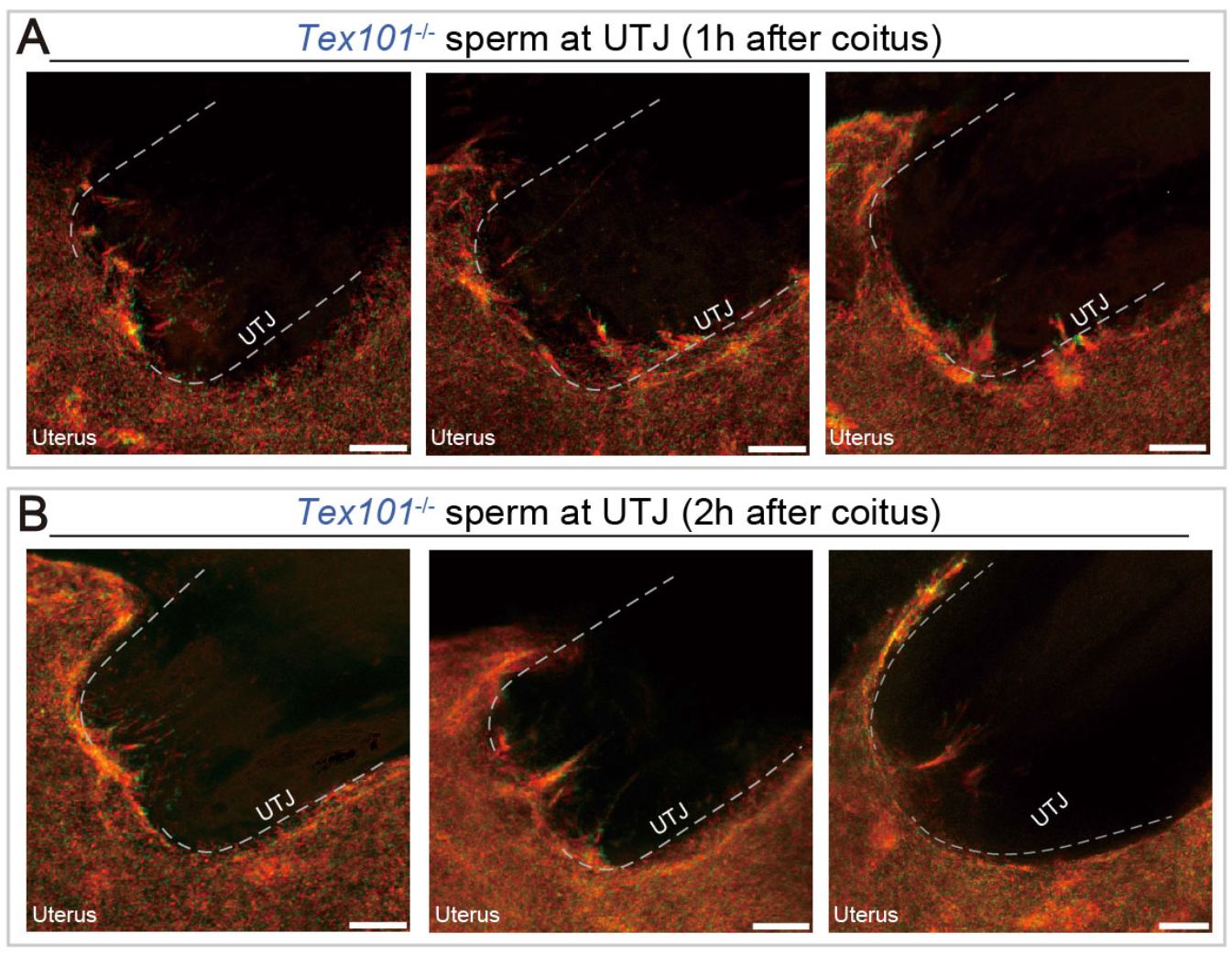
*Tex101*^−/−^ sperm behavior at UTJ. (A) *Tex101*^−/−^ sperm behavior at UTJ 1h after coitus. (B) *Tex101*^−/−^ sperm behavior at UTJ 2h after coitus. Scale bars, 200μm.

**S3 Fig.**
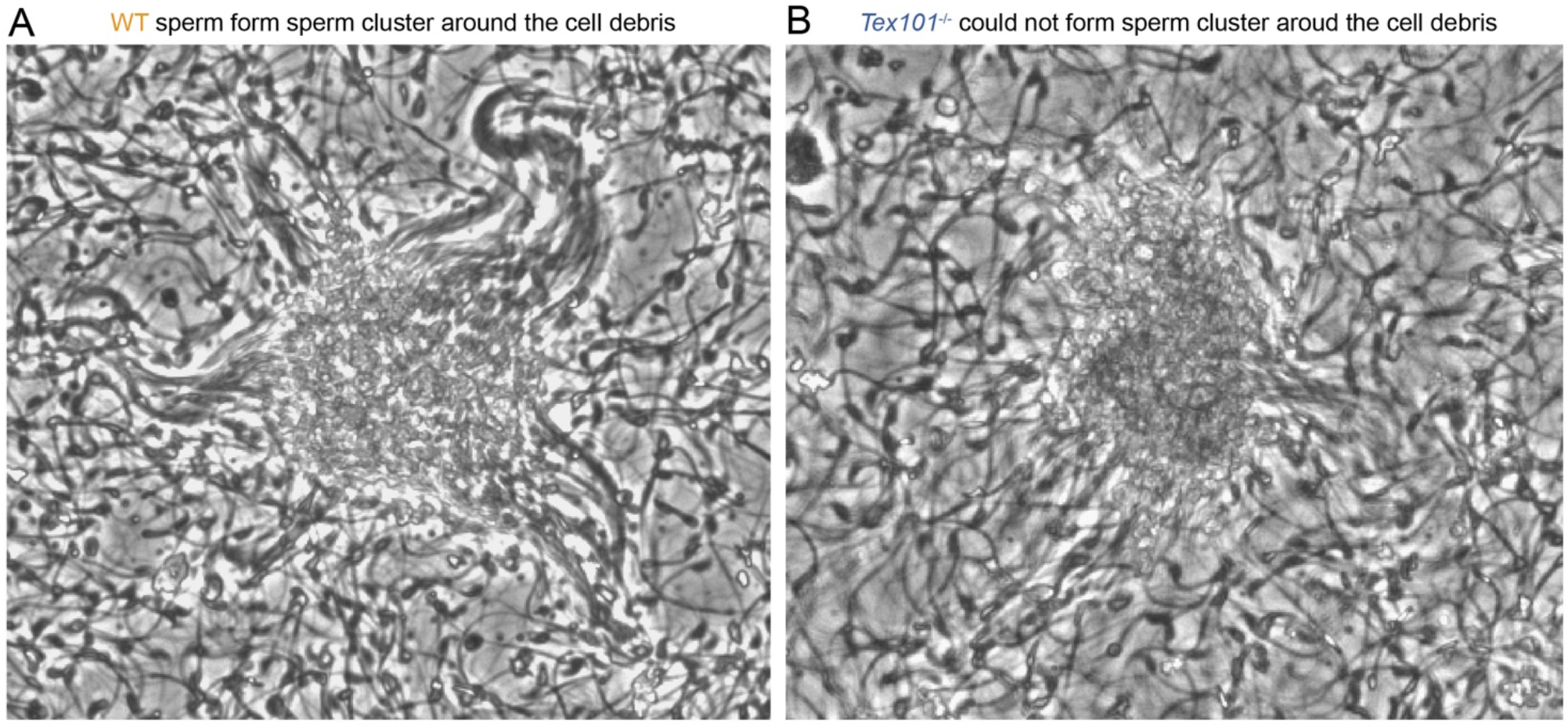
Synchronized sperm tail bending in sperm cluster. (A) Synchronized WT sperm tail bending in sperm cluster 1.5h after coitus. (B) *Tex101*^−/−^ sperm could not form sperm cluster and form the synchronized tail bending.

**S1 Movie.**
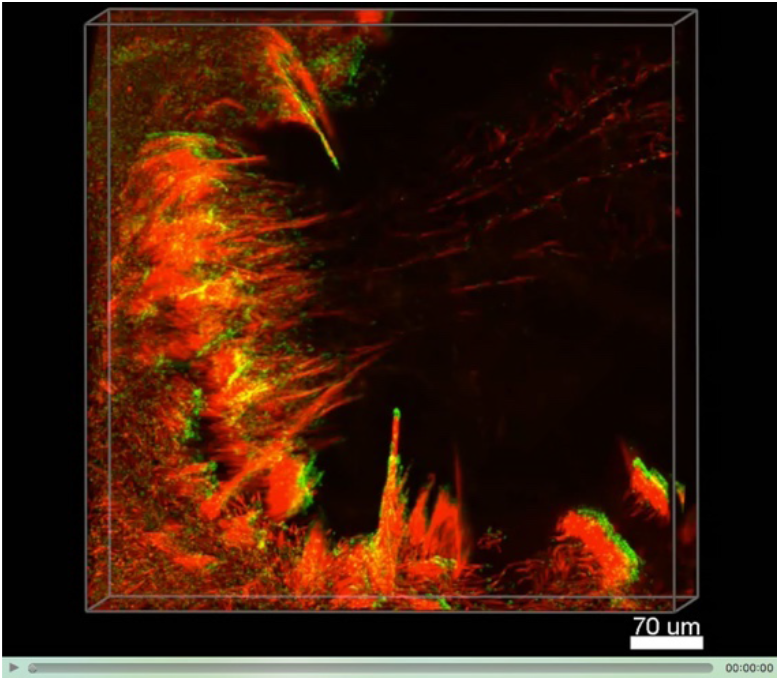
WT Sperm behavior at UTJ 1h after coitus.

**S2 Movie.**
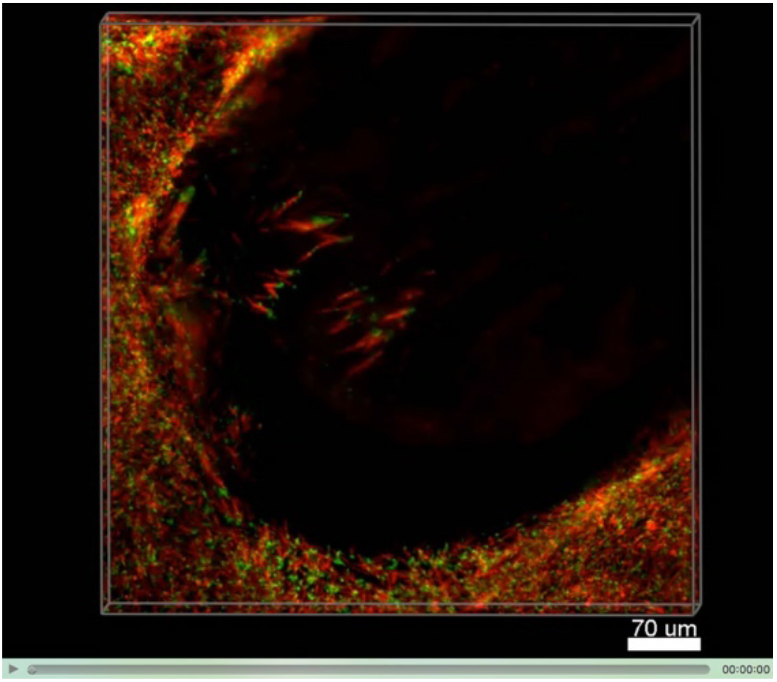
*Tex101*^−/−^ Sperm behavior at UTJ 1h after coitus.

**S3 Movie.**
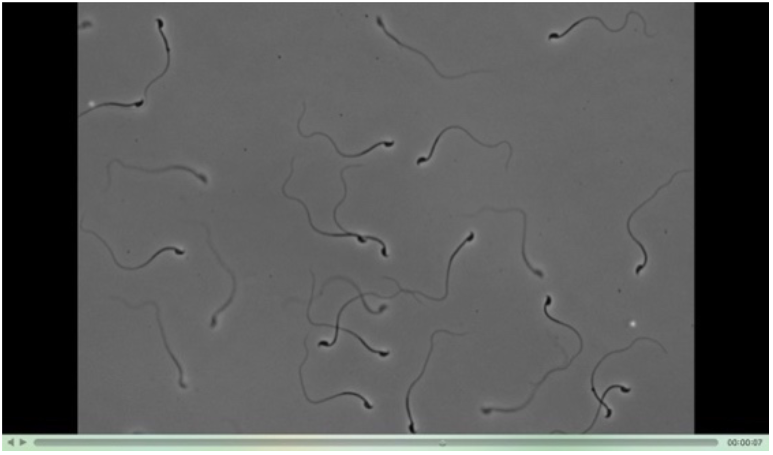
WT Sperm bending pattern in viscoelastic medium 1.5h after coitus.

**S4 Movie.**
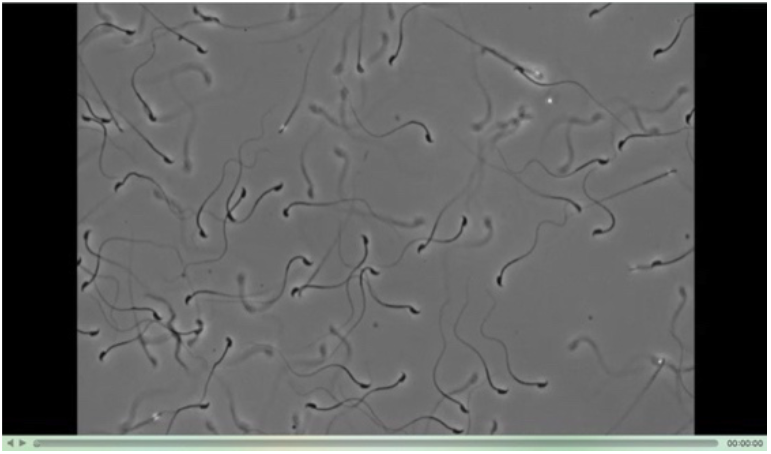
*Tex101*^−/−^ Sperm bending pattern in viscoelastic medium 1.5h after coitus.

**S5 Movie.**
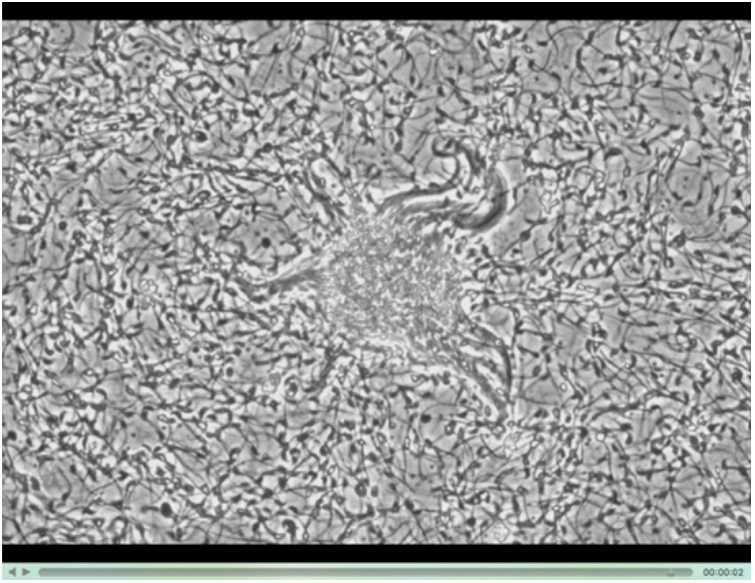
Synchronized WT sperm tail bending around cell debris 1.5h after coitus.

**S6 Movie.**
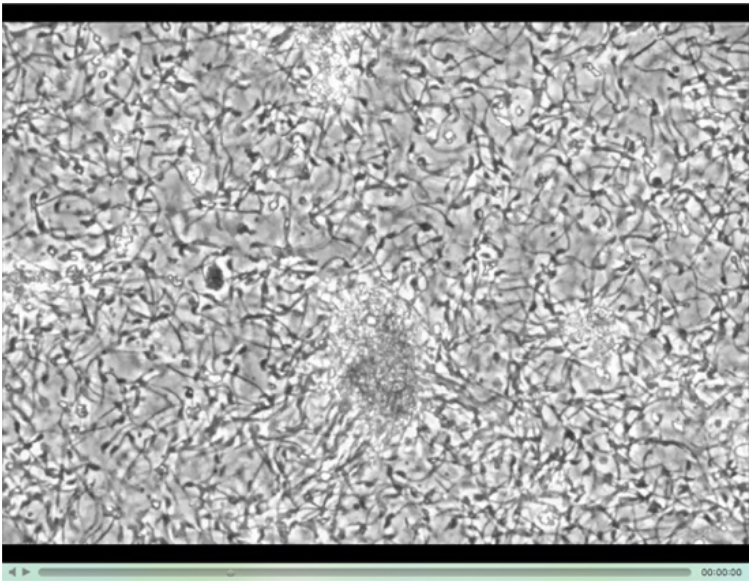
*Tex101*^−/−^ sperm with no synchronized tail bending around cell debris 1.5h after coitus.

